# Inducing a mental context for associative memory formation with real-time fMRI neurofeedback

**DOI:** 10.1101/2022.02.16.480602

**Authors:** Silvy H.P. Collin, Philip van den Broek, Tim van Mourik, Peter Desain, Christian F. Doeller

## Abstract

Memory, one of the hallmarks of human cognition, can be modified when humans voluntarily modulate single neuron or neural population activity using neurofeedback. However, it is currently unknown whether memory is facilitated or impaired after such neural perturbation. In this study, participants memorized objects while we trained them with abstract neurofeedback to modulate their brain activity patterns in the ventral visual stream to represent either faces or houses. The neurofeedback created an implicit face or house context in the brain while memorizing the objects. The results revealed that participants created associations between each memorized object and its implicit context solely due to the neurofeedback manipulation. Our findings shed light onto how memory formation can be influenced by synthetic memory tags with neurofeedback and advance our understanding of mnemonic processing.

## Introduction

Humans can be trained to voluntarily modulate neural activity in various brain regions, which has been shown to influence behavior ^1,2,11–13,3–10^. Animal studies using optogenetics have demonstrated that modulating neurons can influence memory ^14,15^ Furthermore, human intracranial studies have shown that behavior is influenced when participants voluntarily modulate activity of single neurons in the medial temporal lobe (MTL) ^16^, or the functional Magnetic Resonance Imaging (fMRI) signal using neurofeedback ^17–20^. However, it remains unknown if and how neurofeedback can influence the integration of memories.

Memory has been shown to depend on the context in which to-be-remembered material was studied ^21,22^ Training participants to voluntarily modulate their neural activity in a particular way (e.g. by providing neurofeedback based on what information was represented in the brain) would lead to better control over the mental context of the participants during neural modulation. Additionally, memory performance has been shown to be vulnerable to interference from other (related) memories ^23–25^. Possibly, neural modulation at specific times (as used by e.g. Suthana et al, 2012 and Jacobs et al, 2016) might have induced interfering memories coming to mind which then influenced memory performance of to-be-remembered information.

In this experiment, we investigated whether voluntary modulation of neural activity in the ventral visual stream (by neurofeedback) can facilitate associative memory using non-invasive fMRI in humans. The experiment (see Fig 1) started with an fMRI-session that included a training block, and two neurofeedback blocks. The training block contained pictures of faces and houses, and was used to train a classifier to distinguish brain activity patterns evoked by those stimulus categories. During the two neurofeedback blocks, each trial started with an image of an object, and was then followed by an abstract presentation of the face (one block) or house (the other block) classifier evidence (see Fig 1B). The order of these two neurofeedback blocks was unknown to the participants. This manipulation trained participants to voluntarily modulate their neural activity, and created an implicit ‘face mental context’ in one block, and ‘house mental context’ in the other block. The MRI-session was followed by a behavioral memory session that included a neurofeedback context test in which memory was tested for the artificial context created by the neurofeedback, and subsequently an associative learning task. The neurofeedback context test was a two-alternative forced choice task during which the participants were presented with the same objects as during the neurofeedback blocks, and had to indicate for each object whether it belongs to faces or to houses. During the associative learning task, each of the objects from the neurofeedback blocks was associated with a specific exemplar (face or house). Half of the objects were associated with a specific exemplar (face/house) of the same category as they received neurofeedback on in the fMRI-session, and the other half of the objects were associated with a specific exemplar from the other category as they received neurofeedback on. We tested their memory of these associations for the category (face or house) and the specific exemplar.

**Figure 1:**
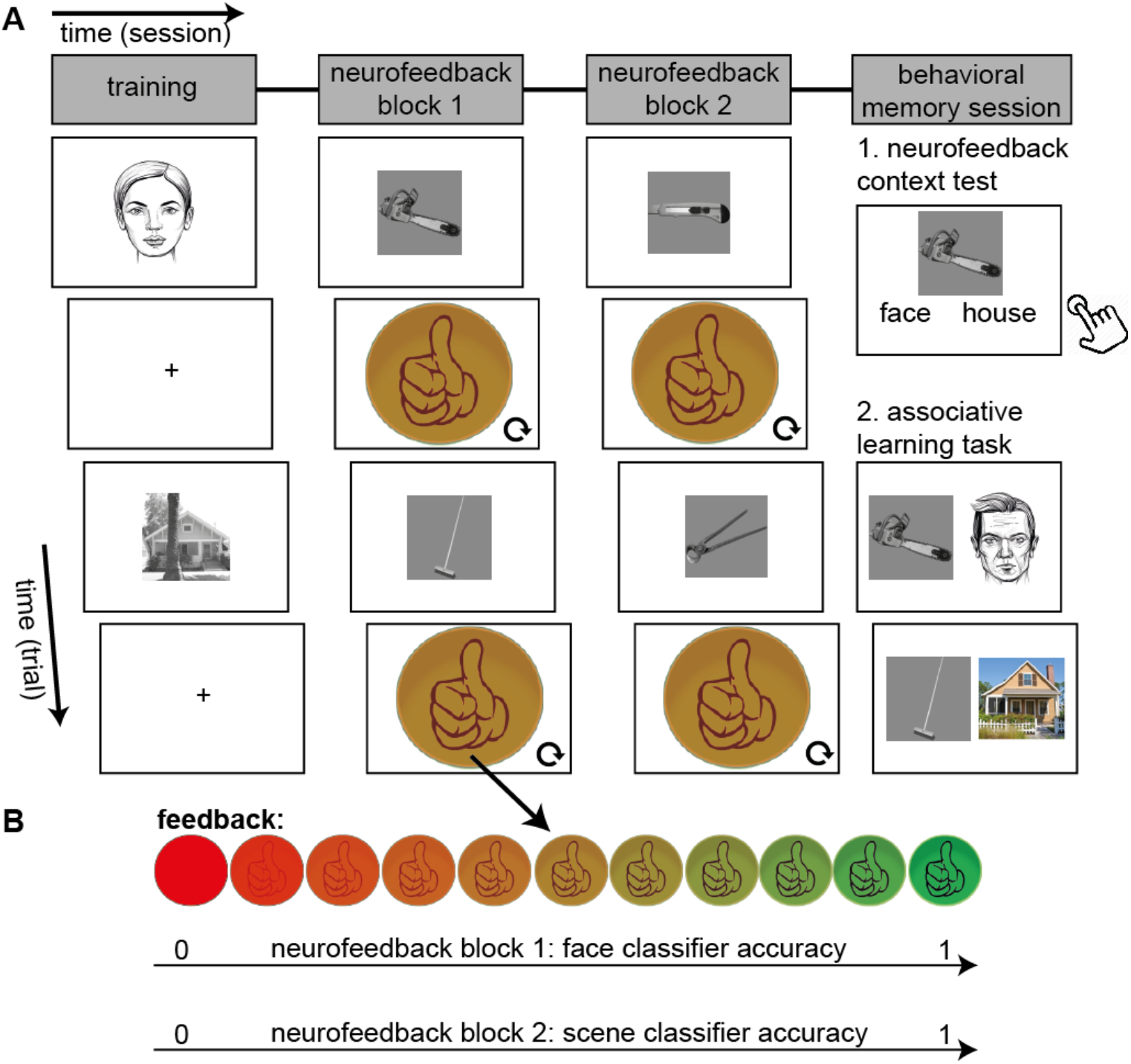
Structure of experimental sessions and neurofeedback. **(A) Overview of the experiment.** The experiment started with an MRI-session that included a training block, and two neurofeedback blocks. The training block contained pictures of faces and of houses, and was used to train a classifier on brain activity patterns evoked by faces and houses. During the two neurofeedback blocks, each trial (24 s duration) started with an image of an object (for 2 s), and was then followed by an abstract presentation of the face (one block) or house (the other block) classifier accuracy (see panel B). The MRI-session was followed by a behavioral memory session that included a neurofeedback context test in which memory was tested for the artificial context created by the neurofeedback. The neurofeedback context test was a two-alternative forced choice task during which the participants were presented with the same objects as during the neurofeedback blocks, and had to indicate for each object whether it belongs to faces or to houses (with a button press). Subsequently participants were presented with an associative learning task. During the associative learning task, each of the 32 objects from the neurofeedback blocks was associated with either a specific face or a specific house. Half of the objects were associated with a specific exemplar (face/house) of the same category as they received neurofeedback on in the MRI-session, and the other half of the objects were associated with a specific exemplar from the other category as they received neurofeedback on. After learning of these pairs, they filled in a Vividness of Visual Imagery Questionnaire, followed by a memory test. Here, it was tested for each object whether they remembered which category it was associated with, and subsequently, with which specific exemplar it was associated. Note that actual face stimuli as used in the experiment are replaced by face drawings in the figure for privacy purposes. **(B) Neurofeedback.** For each trial, the neurofeedback started with an orange circle in which a hand with a thumb up was presented. Based on the classifier accuracy, the alpha-level of the image was adapted, and could therefore change into a red circle (low classifier accuracy) or a green circle with a clearly visible thumb (high classifier accuracy). In one block, the classifier accuracy corresponded to the face category, and in the other block to the house category. The order of the two blocks were counterbalanced across participants.

We hypothesized that it is possible to create a category-specific mental context in higher order visual regions by neurofeedback. We predicted that this mental context would be accompanied by significant decoding evidence of the associated category after participants were exposed to the neurofeedback for a sufficient time, i.e. at the end of each of the blocks. We predicted that the (during neurofeedback) studied objects will become associated with the specific context (face or house) in which they are being encoded ^22,26^. Furthermore, we predict that subsequent use of the same objects in an associative memory task will facilitate associative memory for the category, and furthermore modulate memory for the specific associations, either by facilitating associative learning ^11^ or by interfering with associative learning ^13^.

## Results

### Real-time decoding from higher order visual regions

Before conducting a study using classifier evidence from higher order visual regions (i.e. parahippocampal gyrus and fusiform gyrus, see Methods) as neurofeedback, it is important to establish a technical set-up (see Fig 2) that would reliably classify an associated item (when it is not presented on the screen) at a single subject, volume-by-volume level. This was verified in a separate pilot experiment (see Supplementary Information, and Supplementary Fig. 1 and 2). Based on these pilot results, we predicted to find increased classifier evidence for the correct category in the neurofeedback trials of the main experiment from Time-in-trial (i.e. MR volume) 4 to 9 (see Fig 3), which is why we focused our MR analyses on the neurofeedback trials in the main experiment on these time points.

**Figure 2:**
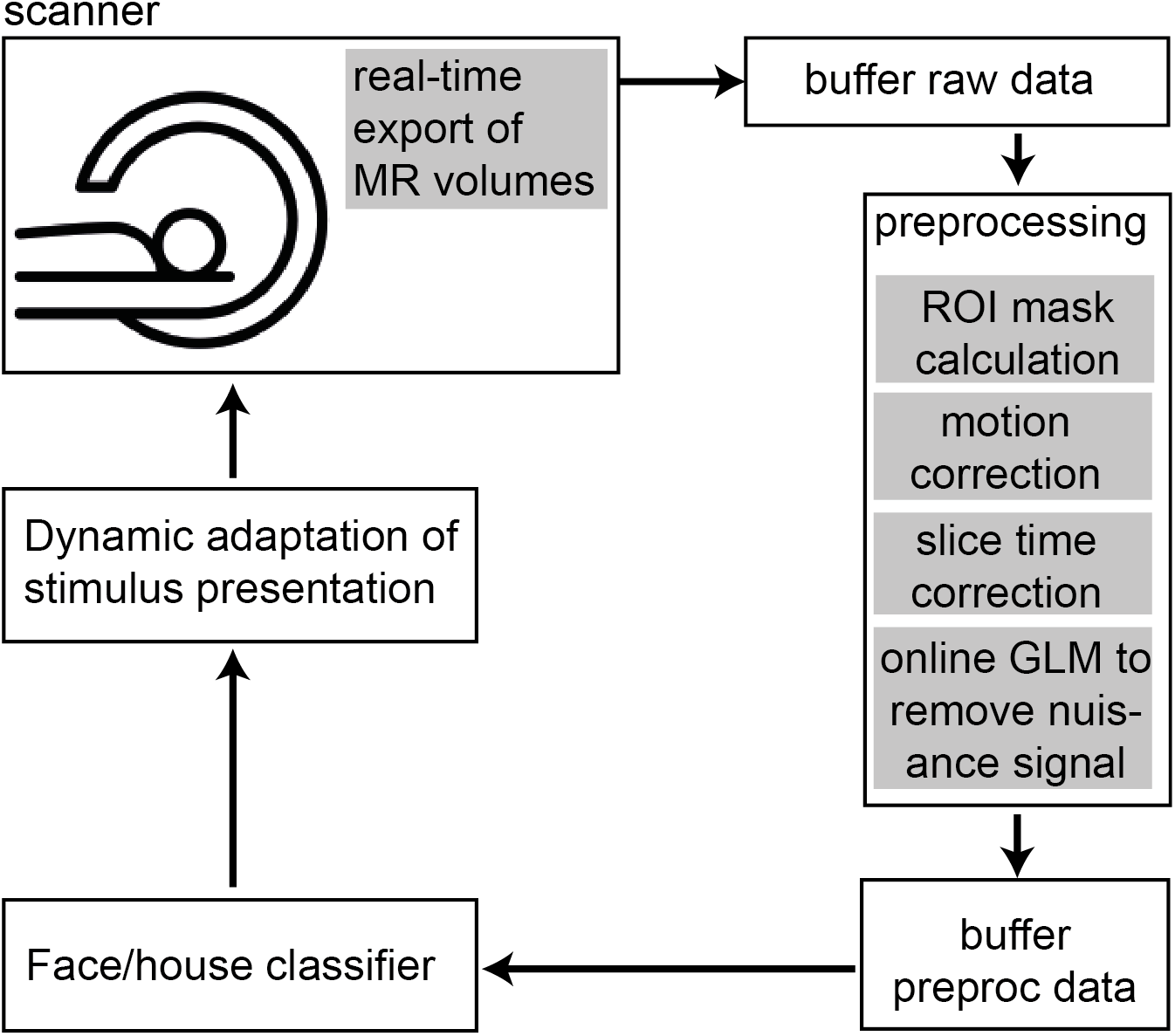
Technical set-up. The technical set-up used for real-time fMRI as used by ^27^ The data was exported in real-time into a FieldTrip buffer, and then immediately preprocessed. A structurally defined ROI mask was calculated. The mask and preprocessed data were imported into BrainStream for online decoding analysis. See methods for more details.

**Figure 3:**
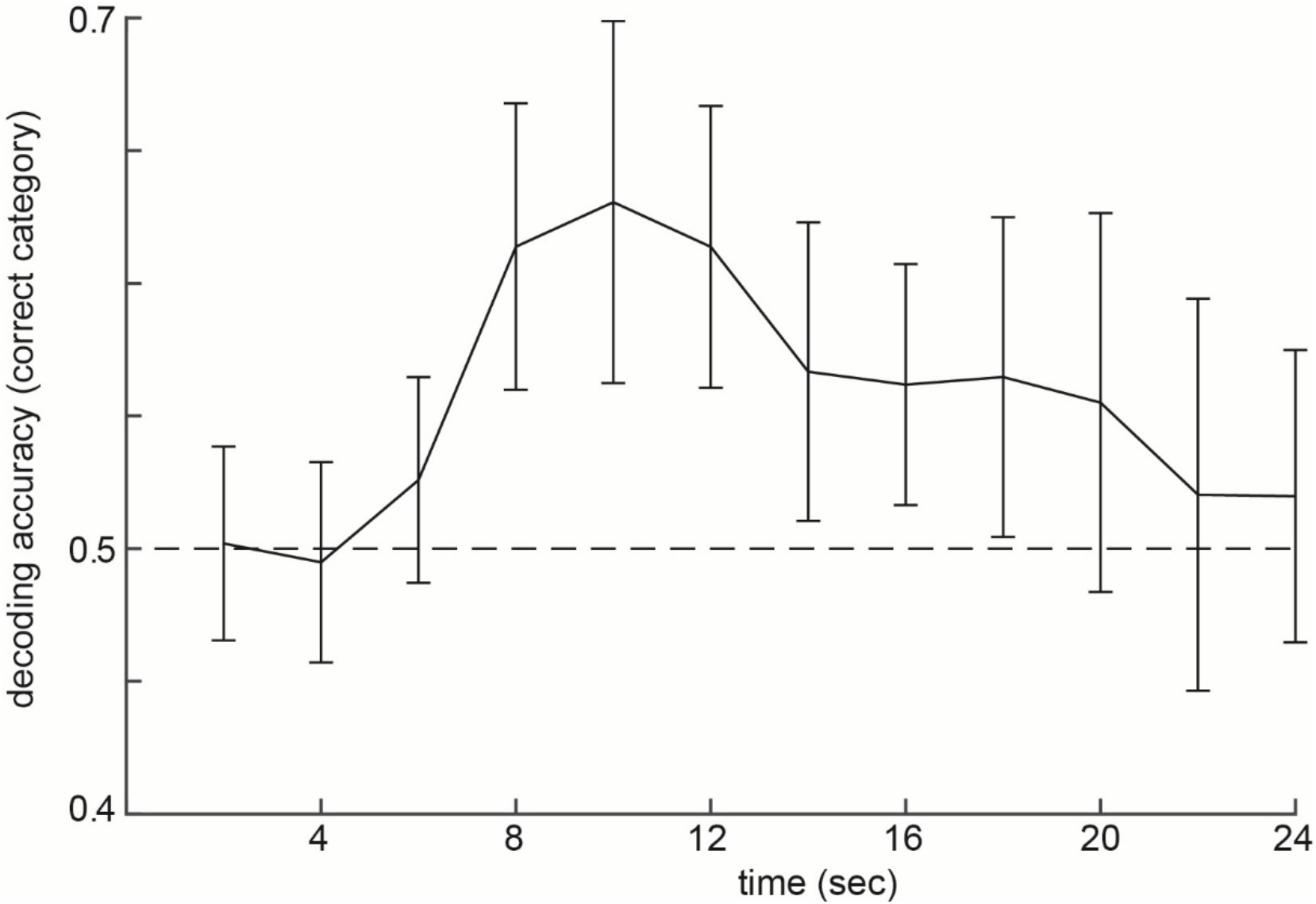
Group average of the decoding evidence during the test phase of the pilot experiment. Decoding evidence (group average ± S.E.M.) of the correct category (0.5 is chance level, indicated by the dashed line). The object is presented on the screen the first 2 s.

### Neurofeedback

The neurofeedback manipulation indeed showed above chance classifier evidence for the correct category (i.e., faces in one block and houses in the other block, based on what was being trained on with the neurofeedback), even though the participants were not told when they had to imagine faces and when they had to imagine scenes. As expected, this was only the case at the end of each neurofeedback block (overall mean: T (1,18) = 2.203, p = 0.041; run 3: T (1,18) = 2.456, p = 0.024; see figure 4).

**Figure 4:**
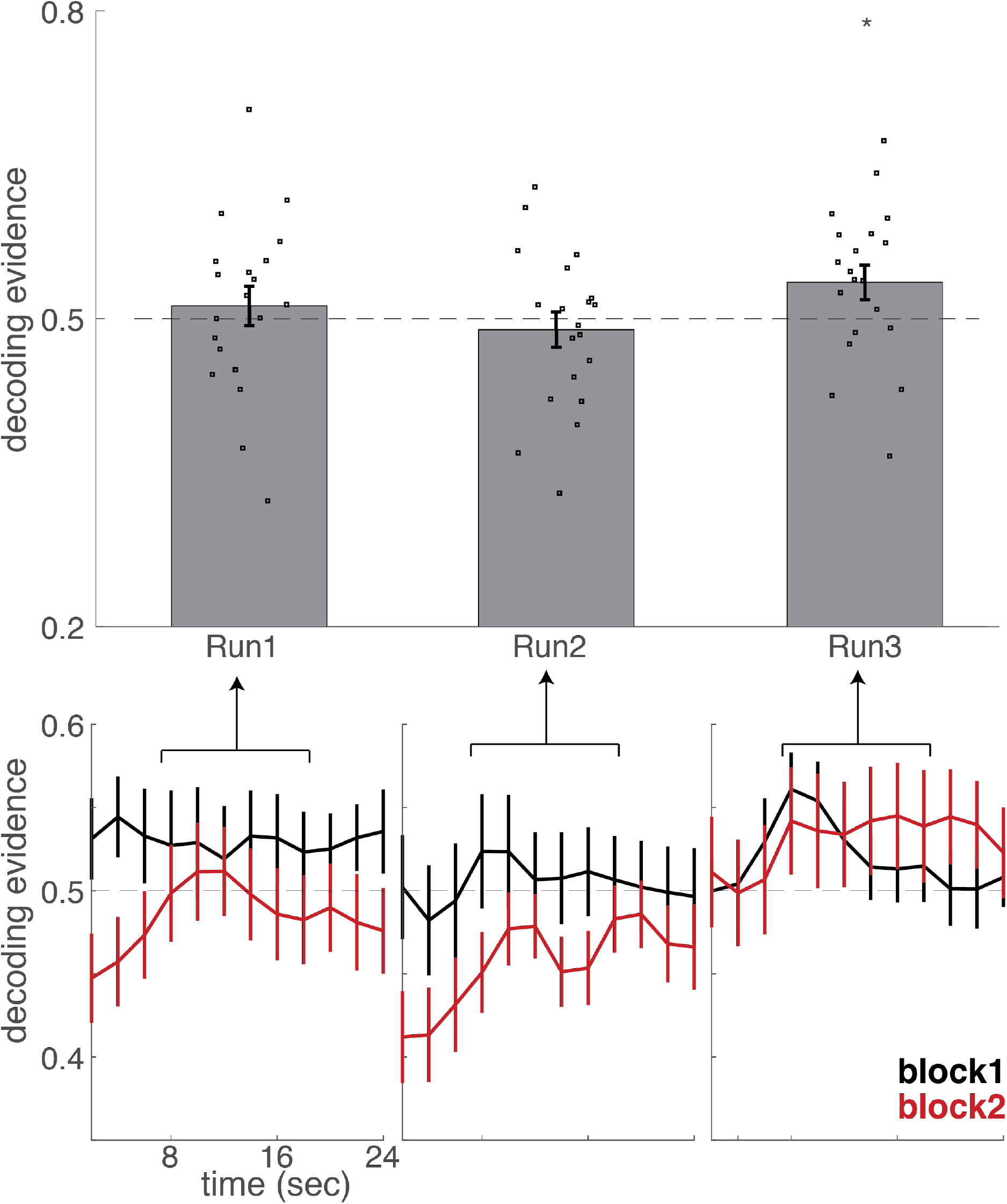
Decoding evidence. Decoding evidence of the correct category during neurofeedback, separately for run 1, 2 and 3 (averaged across all trials of that run), plotted for neurofeedback block 1 and 2 separately. 0.5 is chance level, indicated by the dashed line. Error bars represent S.E.M. The object is presented on the screen the first 2 s, the neurofeedback presentation (see Fig. 1B) is presented in the remaining 22 s. One subject was an outlier and therefore excluded (i.e. more than 2.5 S.D. from group mean). The below plots show the entire time line, and the above plot the mean of run 1, 2 and 3 of the critical time points (based on the pilot experiment, time point 4 to 9, dots represent single subject data). Overall mean: T (1,18) = 2.203, p = 0.041; run 1: T (1,18) = 1.066, p = 0.301; run 2: T (1,18) = - 0.136, p = 0.894; run 3: T (1,18) = 2.456, p = 0.024.

### Neurofeedback facilitated associative memory

Strikingly, the neurofeedback context test revealed that participants indeed associated the memorized objects with their corresponding neurofeedback context. Thus, half of the memorized objects became associated with the face-context induced by neurofeedback, and the other half of the memorized objects became associated with the house-context induced by neurofeedback. It was counterbalanced across participants which object became associated by neurofeedback with which category (T (1,18) = 3.007, P = 0.008, see figure 5).

**Figure 5:**
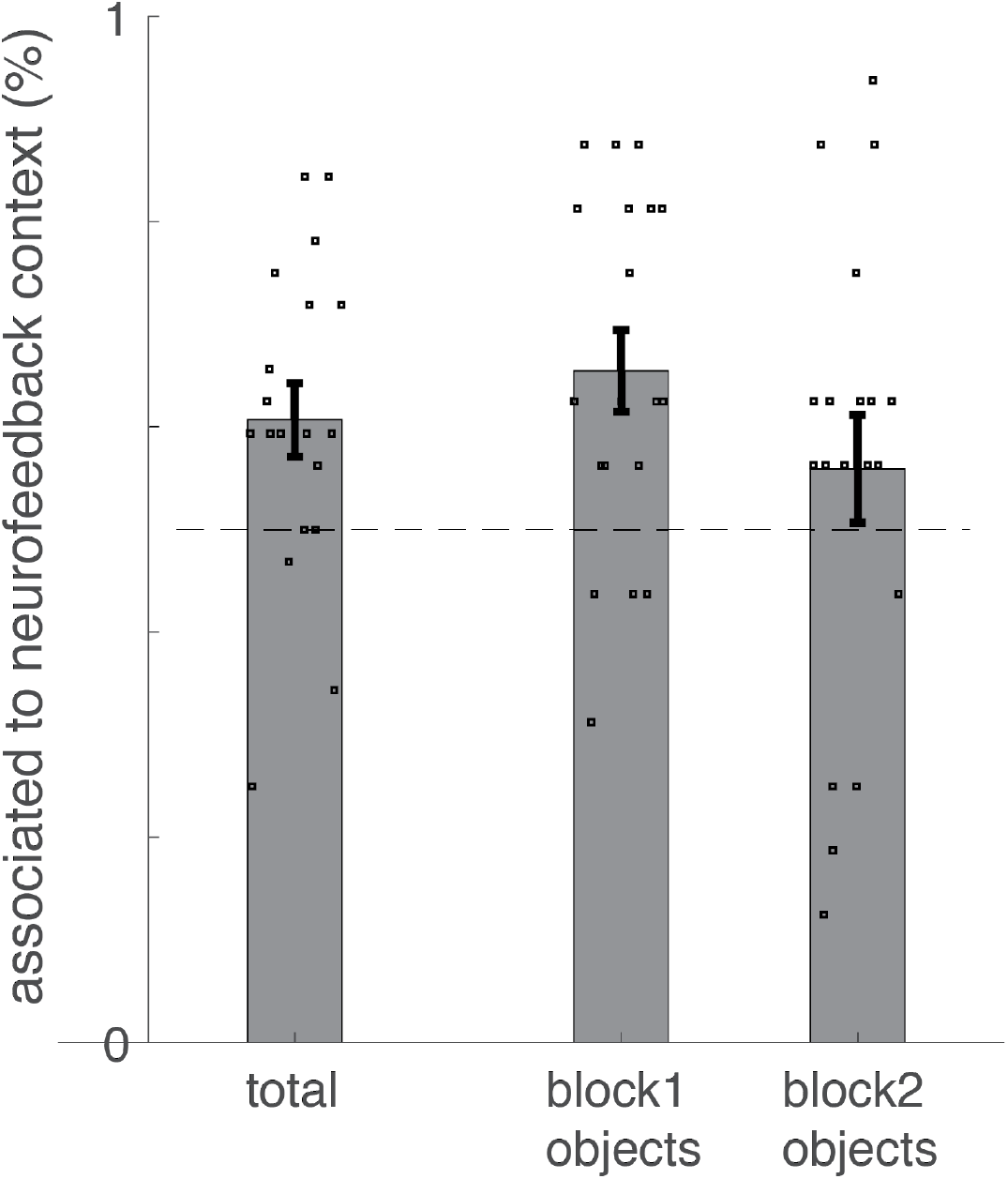
Results of the neurofeedback context test. Average (± S.E.M.) percentage of correct responses during the neurofeedback context test (overlaid with individual responses). From left to right: average across all objects, average across those objects studied during neurofeedback block 1, average across those objects studied during neurofeedback block 2. One subject was an outlier and therefore excluded (i.e. more than 2.5 S.D. from group mean). Total: T (1,18) = 3.007, P = 0.008.

During the subsequent associative learning task (see figure 6), participants had to memorize pairs of images, consisting of the same objects as used in the neurofeedback blocks but now explicitly associated with a novel face or house. Each object was either associated with the same or the other category compared to the neurofeedback context. We predicted that it would modulate associative memory when an object would be associated with the same context as the initial neurofeedback context. However, there was no difference between same category pairs and other category pairs for category memory (T (1,19) = 0.448, p = 0.659) nor for exemplar memory (T (1,19) = - 1.406, p = 0.176).

**Figure 6:**
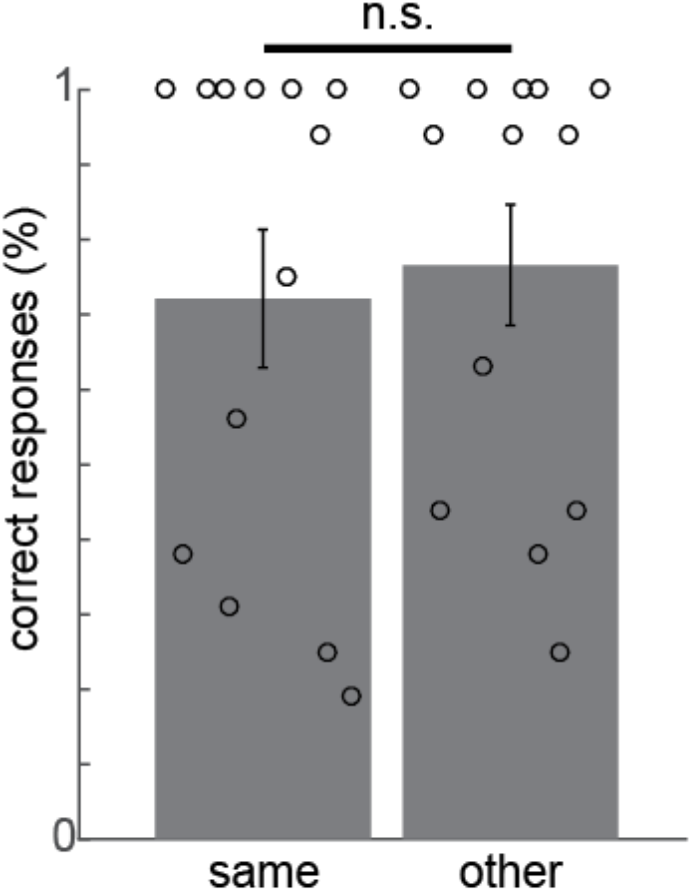
Results of the subsequent associative learning task. Average (± S.E.M.) percentage of correct responses during the subsequent associative learning task, plotted separately for same category pairs and other category pairs (overlaid with individual responses). There was no significant difference between same and other category pairs’ memory.

## Discussion

In this study, participants memorized objects while we trained them with neurofeedback to modulate their brain activity patterns in ventral visual stream regions to simultaneously represent either faces or houses. The results revealed that the neurofeedback manipulation led to the creation of a category specific mental context in the ventral visual stream. Crucially, this caused participants to associate the memorized objects with its specific mental context, either face of house, solely based on abstract neurofeedback.

We successfully trained participants to modulate across-voxel neural representations in ventral visual stream regions using fMRI neurofeedback in such a way that it would represent faces in one half of the experiment and houses in the other half of the experiment, creating an implicit face and house context while having to memorize objects. These face and house representations emerged solely by the continuously adapted, abstract presentation of individual face-house classifier output as feedback to the participants. Thus, crucially, without explicitly notifying them when to think about what category. This is in line with findings from earlier studies in which it has been demonstrated that participants can learn to voluntarily control their across-voxel neural patterns by neurofeedback ^4,17–19,27^. Importantly, our results reveal the possibility of solely using neurofeedback to train participants to associate (more complex) stimulus categories in higher order regions to objects, which extends the neurofeedback study from Amano and colleagues ^19^ in which an association between a color and a grating orientation was created in the early visual cortex by training participants to modulate across-voxel neural patterns in their early visual cortex to represent a specific color while presenting a specific orientation on the screen.

Additionally, we tested how the neurofeedback influenced associative learning. The results revealed that the mental context solely created by neurofeedback caused associative learning, i.e. objects became associated to their corresponding context as shown by a post-scanning memory test. Memories are contextualized, and remembering something will lead to recovery of the context in which that memory was formed ^28^. Empirical research showing the importance of context for memory ^29–31^ dates back to the famous Godden and Baddeley study showing the influence of a land vs underwater context on memorizing words ^32^. Less is known about how malleable the neural underpinnings of a memory context is ^29,33^. These results demonstrate the malleability of a mental context by showing that it is possible to voluntarily modulate the neural representations of a mental context to represent a specific image category (in our case faces or houses) while another task is being performed (in our case learning a list of objects). Furthermore, we provide evidence for the notion that this mental (face/house) context implicitly influences (object) memories that are being formed in this context.

Intracranial studies in humans revealed that memory performance after modulating MTL neurons could lead to both memory facilitation ^11^ as well as memory impairment ^13^. However, in contrast to what we expected based on earlier literature, we did not observe an effect of the neurofeedback manipulation on a subsequent associative learning task with novel images. It could possibly be due to participants approaching ceiling level performance in this task that we did not confirm our prediction. Future research would be necessary to draw more firm conclusions about this outstanding question.

Our study investigated how paired associate learning is influenced by an implicit artificial context that was created by voluntary modulation of neural populations. The results open interesting avenues for future research investigating how to influence memory integration and inference with neurofeedback, and enhances our understanding of mnemonic coding in the brain.

## Author contribution

SHPC, PD and CFD designed the experiment; SHPC, PvdB, and TvM were involved in the initial pilot and implementing the technical set-up; SHPC collected and analyzed the data; all authors contributed to writing the manuscript.

## Methods

### Participants

Thirty students from the Radboud University campus in Nijmegen participated in this study. All participants were right-handed and had normal or corrected-to-normal vision. Ten participants had to be excluded to do technical problems, excessive head motion, or an incomplete dataset. Thus, the final group of participants contained twenty students (nine males, aged 20-44 years, mean age 27.2). All participants gave written informed consent. The study was approved by the local ethics committee (CMO, Arnhem/Nijmegen).

### Task design

The experiment consisted of a training block, two neurofeedback blocks of 3 runs each, a neurofeedback context test, and an associative learning task (see Fig. 1). The tasks were presented using Brainstream/Psychtoolbox (training block and neurofeedback blocks) or Presentation software (neurofeedback context test and associative learning task; Neurobehavioral Systems, version 16.4).

### Training block

The training block was used to train a classifier on brain activity patterns evoked by faces and houses. The training block consisted of 28 blocks in total, interleaved blocks with images of faces and blocks with images of scenes. Each block lasted for 30 seconds and was followed by a fixation cross which was presented for 12 seconds. Each block consisted of 14 unique pictures (i.e. each block had a different set of pictures), each picture was presented for 2 seconds, and, additionally, the first picture of the block was repeated at a random position within that block. Participants had to press a button when they saw the first picture being repeated. They were asked to maintain attending the images throughout the entire block.

### Neurofeedback blocks

During the two neurofeedback blocks, participants received neurofeedback based on the evidence of the classifier after being presented with an image of an object. The object was presented during the first 2 s, and the neurofeedback was being presented for the subsequent 22 s. Each neurofeedback block had 16 unique objects that were all presented three times (all 16 objects are shown once in a random order before the first object is being repeated), which led to a total of 32 unique objects. The neurofeedback was presented in an abstract fashion; by manipulating the color of a circle, and the visibility of a hand with the thumb pointing up (see Fig. 1B). If the decoding evidence of the correct category was above chance level (i.e. above 50 %), then the alpha level of the green neurofeedback image (i.e. the image with the green circle that includes an image of a hand with the thumb pointing up) increased with 0.05. If the decoding evidence of the correct category was below chance level, the alpha level of the green neurofeedback image decreased with 0.05 (making the circle appear more red, and the hand less visible). For one block the correct category was ‘face’, and for the other block the correct category was ‘house’, with the order counterbalanced across participants. This leads to 16 objects only being presented during neurofeedback training of the face category, and the other 16 objects being presented only during neurofeedback training of the house category.

### Neurofeedback instruction

Participants were instructed that the neurofeedback image will change based on their brain activity, and that they can influence this by thinking about either faces or houses. They were *not* told which actual category they have to think about in which block, and were told to figure this out based on the neurofeedback that they receive. Furthermore, they were told to try to vividly imagine images of that category during the neurofeedback blocks, and that they receive 3 euros extra monetary compensation with good performance. The expected 5-6 sec delay in the neurofeedback (due to the BOLD response) was explained to them as well. They are also told to remember which objects are shown.

### Neurofeedback context test

After the real-time fMRI session, participants were placed behind a computer screen, and conducted a neurofeedback context test. During this (self-paced) test, participants were presented with the 32 objects from the neurofeedback blocks, one at the time. For each object, they were asked to which category this object belonged (face or house).

### Associative learning task

Afterwards participants performed an associative learning task. They were presented with 32 pairs. Each pair consisted of one object and one face, or one object and one house. The faces and houses used were different from the ones used in the training block. The objects were the same as the ones used in the neurofeedback blocks. Half of the objects were associated with an exemplar (face or house) from the same category as the category on which they received neurofeedback for this object (referred to as same category pairs). The other half of the objects were associated with an exemplar (face or house) from the other category as the category on which they received neurofeedback for this object (referred to as other category pairs). There were twelve repetitions of each pair (in random order) leading to 384 trials in total, with each trial showing the images for 2.8 s and a fixation cross for 0.2 s. After a short break during which participants filled in a Vividness of Visual Imagery Questionnaire, they were tested for their memory for the 32 pairs by asking them for each of the 32 objects whether it the object belongs to a face or to a house, and subsequently to which specific face or house this object belongs. Responses were given with button presses. For the second question they received 16 possible answer options (i.e. the 16 exemplars, 8 faces and 8 houses, that were associated with one of the objects from neurofeedback block 1 were the answer options for all neurofeedback block 1 objects, and similar for neurofeedback block 2 answer options). They select their answer with a button press. The objects were presented in a random order.

### Image acquisition

All images were acquired using a 3T Siemens Prisma scanner equipped with a 32-channel head coil (Siemens, Erlangen, Germany). The structural T1-weighted image was acquired using an MPRAGE-grappa sequence with the following parameters: TR = 2300 ms; TE = 3.03 ms; flip angle = 8°; inplane resolution = 256×256 mm; number of slices = 192; acceleration factor PE = 2; voxel resolution = 1 mm^3^, duration = 321 s. The functional images were acquired using a 2D Echo Planar Imaging (EPI) sequence, with the following parameters: voxel size 3.3 × 3.3 × 3 mm, TR = 2000 ms, TE = 30 ms, flip angle = 80 deg, Multi-Band acceleration factor = 2, FOV = 212 × 212 × 105 mm.

### Real-time fMRI analysis

The functional volumes were preprocessed and analyzed in real-time using BrainStream software (see www.brainstream.nu) as was used before, see Ref ^27^, which is a Matlab-based software package developed at the Donders Centre for Cognition (Nijmegen, Netherlands). The toolbox builds on Psychtoolbox combined with an extension (StimBox) for adaptive stimulus presentation, FieldTrip toolboxes for raw and preprocessed data buffers, FSL and SPM8 for MR data analyses, a GUI streamer to access and export the raw MR volumes during acquisition, and the Donders Machine Learning Toolbox for online decoding. See Fig 3 for an overview of the technical set-up.

### Online image preprocessing

Each functional volume was sent to another computer directly after acquisition of the volume, and stored in a Fieldtrip buffer. From this buffer, the scan entered a (matlab based) preprocessing pipeline (BrainStream). This preprocessing pipeline included motion correction (X, Y, Z, pitch, roll, yaw), slice time correction, and an online GLM to remove nuisance signal.

### Online decoding

After preprocessing, the scans entered another Fieldtrip buffer from which they entered the decoding analysis. For the training block the scans were first shifted for 6 secs to account for the hemodynamic delay. Then, the scans were labeled according to their category (face, scene) and used to train a classifier. We used logistic regression in conjunction with an elastic net regularizer, as used in Niazi et al. (2013). Using a coordinate gradient-descent algorithm ^34^, classifier training took only a few minutes to complete (see Ref ^27^ for more details on the classifier). The voxels corresponding to the parahippocampal gyrus and fusiform gyrus were used for classifier training and test (a native space mask was calculated using the inverse normalization parameters to inverse an MNI space mask including parahippocampal gyrus and fusiform gyrus into subject specific space).

## Acknowledgments

CFD’s research is funded by the Kavli Foundation, the Centre of Excellence scheme of the Research Council of Norway – Centre for Biology of Memory and Centre for Neural Computation, The Egil and Pauline Braathen and Fred Kavli Centre for Cortical Microcircuits, the National Infrastructure scheme of the Research Council of Norway – NORBRAIN, the Netherlands Organisation for Scientific Research (NWO-Vidi 452-12-009; NWO-Gravitation 024-001-006; NWO-MaGW 406-14-114; NWO-MaGW 406-15-291) and the European Research Council (ERC-StG RECONTEXT 261177; ERC-CoG GEOCOG 724836). SHPC is supported by the Netherlands Organisation for Scientific Research (NWO-Rubicon grant 446-17-009). The authors would like to thank Alejandro Vicente-Grabovetsky and Sander Bosch for technical support. Correspondence should be addressed to SHPC or CFD.

## Additional information

### Data availability statement

Data and code are available upon request to the authors.

## Supplementary information

### Supplementary pilot experiment

Before conducting a study using classifier evidence as neurofeedback to train participants to integrate memories, it is important to establish a technical set-up that would reliably classify an associated item at a single subject, single trial, volume-by-volume level from our regions of interest, which is the aim of this pilot experiment. We used a basic associative memory design, and adapted that to a real-time fMRI decoding version, in order to test whether we could reliably decode stimulus categories at a single subject, single trial, volume-by-volume level at moments that *not* the being decoded stimulus category but an *associated* item was presented on the screen (see Supplementary Fig. 1 for task design).

**Supplementary Figure 1:**
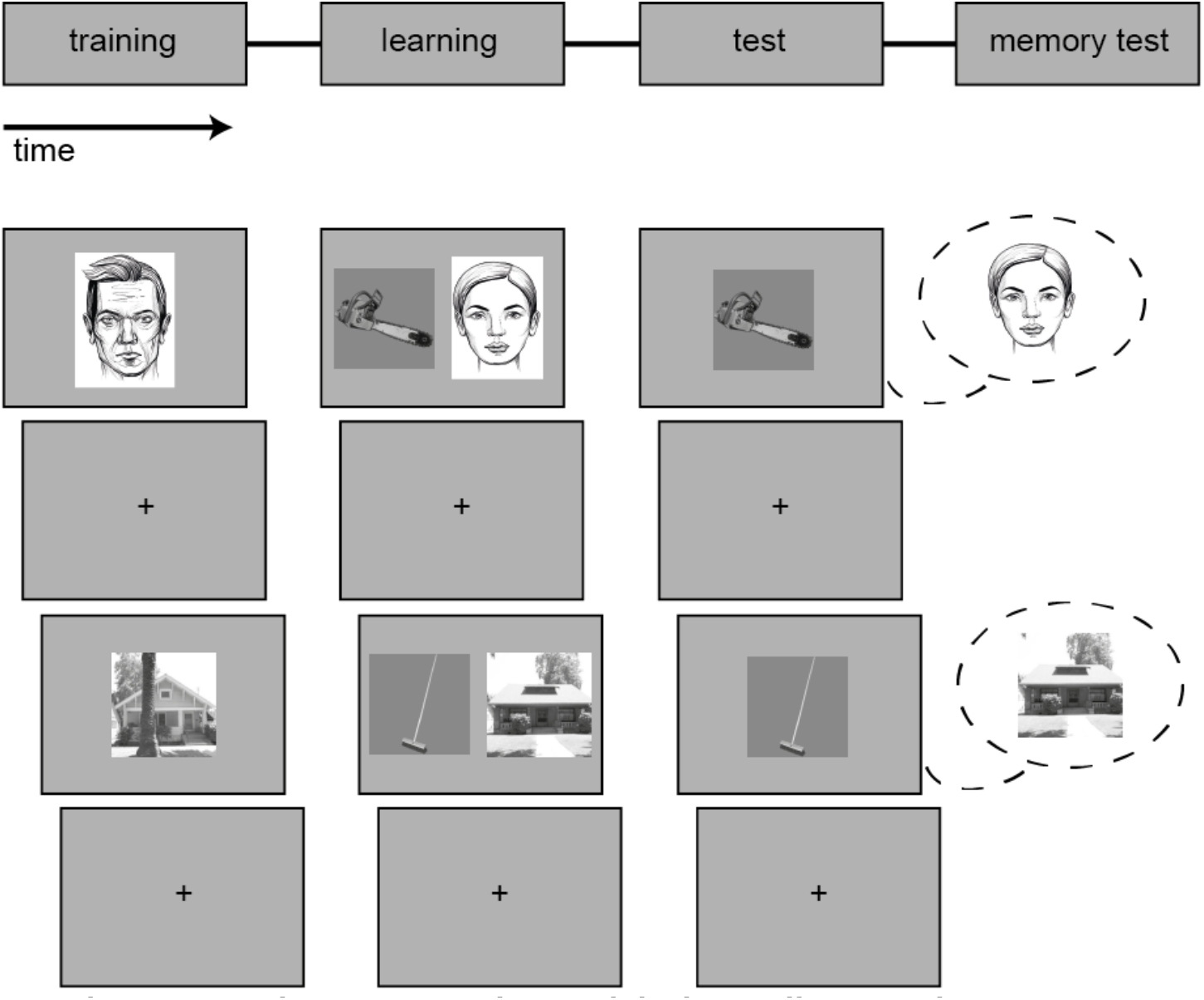
Experimental design – pilot experiment. The pilot experiment consisted of a training phase, learning phase, test phase and memory test. Note that actual face stimuli as used in the experiment are replaced by face drawings in the figure for privacy purposes.

#### Results

We trained a classifier on brain activity patterns evoked by faces and houses. In a learning phase, participants learned four object-face pairs, and four object-house pairs. Subsequently, participants were presented with these eight objects (consecutively in a random order) while we calculated for each MR volume the decoding accuracy of faces and houses from their brain activity patterns (see methods for more details on task and analyses).

The classifier results for scene and face decoding showed variable results across trials/participants. Of the 20 trials per category (5 participants, 4 trials each), scene decoding was above chance level (i.e. 0.5) on 17 out of 20 trials, and face decoding was above chance level on 11 out of 20 trials. When looking at the volume-by-volume results, 36 out of 40 trials in total reached above chance level performance (i.e. above 0.5) at some time during the 24 s trial. The 4 trials that never reached above chance level decoding accuracy were all trials in which the object was associated with a face. A possible explanation could be that the memory performance for the face pairs was lower than the scene pairs at the group level (on average 85% correct for scenes, and 75% correct for faces). Altogether, we conclude that it is feasible with the current set-up to use decoding accuracy as neurofeedback as a tool to train participants to associate stimuli.

**Supplementary Figure 2:**
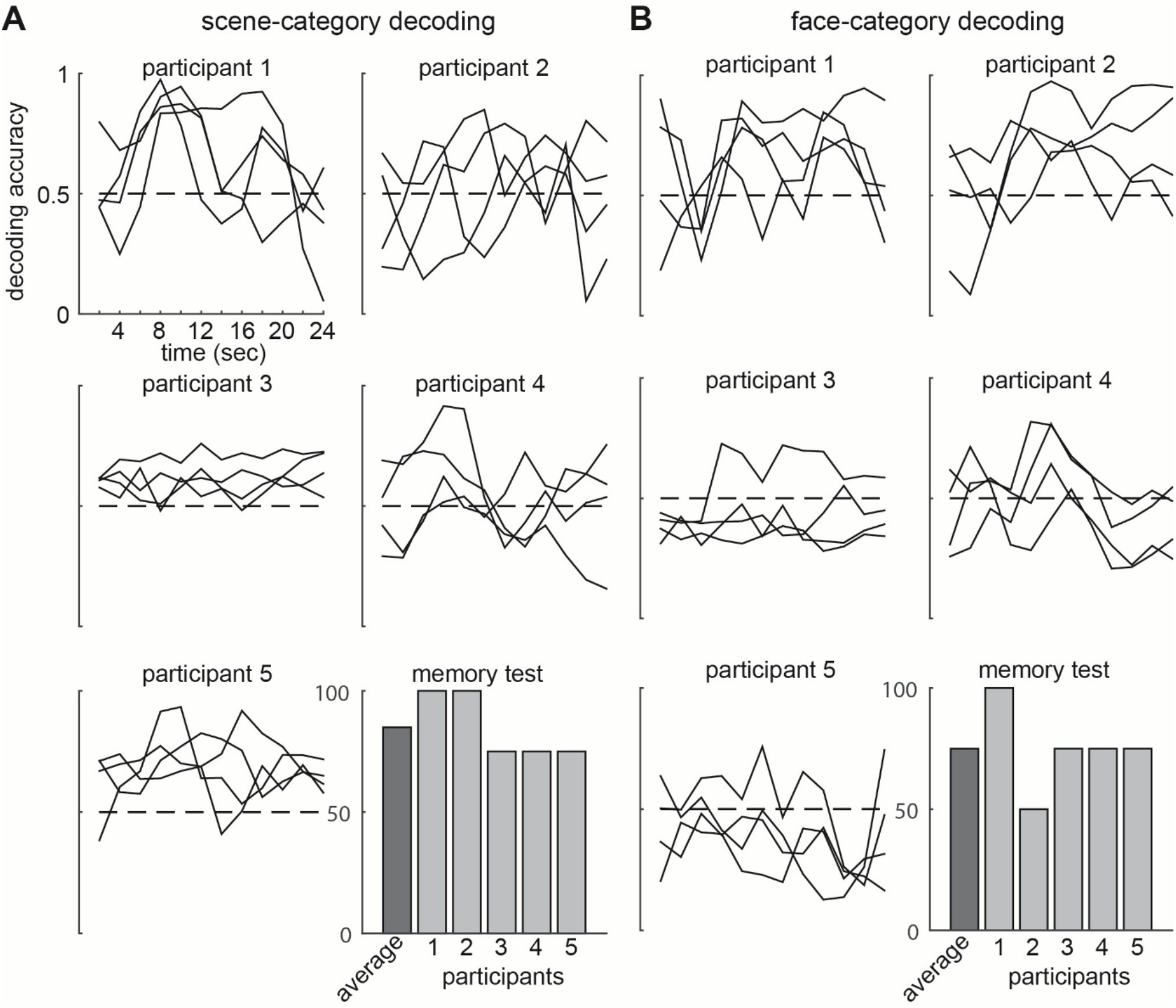
Single subject, single trial, volume-by-volume scene-category (A) and face-category (B) decoding during the pilot experiment. Each plot shows the decoding evidence (of scenes in A, and of faces in B) over time of a single participant for each trial separately (total of 4 trials per participant) during the pilot experiment (as described in supplementary material). During this decoding test phase, the participant was looking at objects that were associated with a scene (in A) or a face (in B) during a prior learning phase. At the bottom right, the memory performance for this category of the group on average (dark bar) as well as each participant separately (light bars) is displayed.

#### Methods

##### Participants

Seven students from the Radboud University campus in Nijmegen participated in this study. All participants were right-handed and had normal or corrected-to-normal vision. Two participants had to be excluded to do technical problems. Thus, five students (one male, aged 19-27 years, mean age 21.2) were included in this pilot study. All participants gave written informed consent. The study was approved by the local ethics committee.

##### Task design

This pilot experiment consisted of four phases; a training phase, a learning phase, a test phase, and a memory test (see Supplementary Fig. 1). The training phase was used to train a classifier on brain activity patterns evoked by faces and scenes. The training phase consisted of 28 blocks in total, interleaved blocks with images of faces and blocks with images of scenes. Each block lasted for 30 seconds and was followed by a fixation cross which was presented for 12 seconds. Each block consisted of 14 unique pictures (i.e. each block had a different set of pictures), each picture was presented for 2 seconds, and, additionally, the first picture of the block was repeated at a random position within that block. Participants had to press a button when they saw the first picture being repeated. They were asked to maintain attending the images throughout the entire block. During the learning phase the participant was required to learn four object-face pairs, and four object-house pairs. Each pair was presented together on the screen 14 times (stimulus duration 1.5 sec, inter-trial interval 0.5 sec). During the test phase, participants were presented with the eight objects from the learning phase. Each object was presented in isolation, with the eight objects presented consecutively in a random order. Each object was presented only once for 24 sec. After the realtime fMRI session, participants were placed behind a computer screen, and conducted a memory test. During this (self-paced) memory test, participants were tested for their memory for the eight pairs. For each object, they were asked to which category this object belonged (face or scene), followed by a certainty rating (scale 1 to 4). Additionally, they were presented with all exemplars from the category used during the learning phase, and asked to identify the correct exemplar which was associated to this object, again followed by a certainty rating (scale 1 to 4).

##### Image acquisition and analyses

Image acquisition, real-time fMRI analysis, online image preprocessing, and online decoding was performed as described in the main methods section.

